# Magic traits, search costs, and the persistence of species in secondary contact

**DOI:** 10.64898/2026.07.23.740407

**Authors:** Jack R. Farley, Darren Irwin

**Affiliations:** Department of Statistics, University of British Columbia, Vancouver, Canada; Department of Zoology, University of British Columbia, Vancouver, Canada

**Keywords:** Hybrid zones, assortative mating, models/simulations, sexual selection, magic traits, search costs

## Abstract

When two populations come into secondary contact, assortative mating can act as a barrier to gene flow. However, when assortative mating is incomplete, associations between preference and cue loci can break down, eroding assortative mating and collapsing species boundaries. Here, we investigate factors that can contribute to the maintenance of differentiated reproductive populations despite gene flow, resulting in either clinal hybrid zones or overlap zones between distinct populations. By simulating secondary contact on a uniform two-dimensional landscape using an individual-based computer model, we examine how the potential outcomes of secondary contact depend on search costs and/or pleiotropy between various combinations of mating cue, mating preference, and hybrid viability traits. We find that search costs can maintain stable mating trait clines, even in the absence of hybrid inviability traits. Pleiotropy between mating cues and hybrid inviability (i.e., “magic cues”) temporarily stabilizes the mating cue cline but fails to maintain mating preference differentiation, since preference is not itself under direct selection. As the preference cline collapses, assortative mating breaks down, and — absent search costs or very strong hybrid inviability — the mating cue cline subsequently collapses as well. Thus, without preference-cue or preference-viability pleiotropy, search costs are essential for maintaining mating trait differentiation. Changes to the genetic architecture of mating traits strongly alter the distribution of phenotypes, affecting whether parental populations can coexist spatially. Small parameter changes can abruptly shift outcomes, highlighting the sensitivity of these systems.

## 1. Introduction

From the songs of warblers to the shapes of orchids to the colours of reef fish, barriers to interbreeding between divergent populations play a major role in maintaining biodiversity (Barreto and McCartney, 2008; Irwin, Bensch, et al., 2001; Schiestl and Schlüter, 2009). Reproductive barriers can take a variety of forms, including behavioural, mechanical, and temporal isolation (Grant and Grant, 1997; Mayr, 1963; Rieseberg and Willis, 2007; West-Eberhard, 1983). However, these barriers are often incomplete and hybrid zones form between closely related taxa (Barton and Hewitt, 1985). Much of our understanding of hybrid zones comes from cline models based on the Fisher-KPP equation (Bazykin, 1969; Fisher, 1937; Kolmogorov et al., 1937). These show that stable clines called “tension zones” can be maintained between two taxa by a balance between the dispersal of parental genotypes into the zone and selection against hybrid genotypes causing a population sink in the middle of the zone (Barton and Hewitt, 1985). The selection against hybrid genotypes is typically attributed to genetic incompatibilities accumulated during allopatry (Dobzhansky, 1937; Muller, 1942), but can also arise through frequency-dependent selection, as with a rare mating-type disadvantage (Bridle et al., 2006; M’Gonigle and FitzJohn, 2010).

Mathematical models show that frequency-dependent selection can maintain stable single-locus clines (Gavrilets, 1997), but assortative mating often requires the divergence of more than one trait. When preference and cue loci are free to recombine, the selection pressure on preference loci can be too weak to maintain linkage disequilibrium (Kirkpatrick and Barton, 1997; Qvarnström, Brommer, et al., 2006), and divergent preferences are vulnerable to homogenization. Because of this, there is interest in mechanisms that can maintain divergent mating preferences. One such mechanism is divergent selection acting on a mating trait, such that the trait’s phenotype and fitness effects are pleiotropic, i.e., influenced by the same genetic loci (Gavrilets, 2004; Jiggins, Emelianov, et al., 2005). Another is search costs, whereby rejecting a potential mate and continuing to search for others comes with a cost in time or energy (M’Gonigle et al., 2012; Real, 1990).

The former mechanism has been termed “magic traits” and can be divided into “magic cues” and “magic preferences” depending on which mating trait the divergent selection is acting (Maan and Seehausen, 2012). Here we require that the divergent selection arises independently of the assortative mating process. This condition is not typically included in the magic trait definition, but is necessary in the context of hybrid zone models to exclude divergent selection resulting from rare mating-type disadvantage. Magic preferences can arise when mating preferences are caused by biases in females’ sensory systems and mating displays targeting those biases then evolve (Basolo, 1990; Dawkins and Guilford, 1996; West-Eberhard, 1984). Magic preferences are thought to be a more potent driver of speciation than magic cues (Maan and See-hausen, 2012); but due to the difficulties in confirming the genetic basis of a preference, their prevalence is debated (Smadja and Butlin, 2011). Mating display traits are naturally more noticeable by researchers, causing magic cues to receive more attention in the literature (Thibert-Plante and Gavrilets, 2013). Initially believed to be rare, recent studies suggest that magic cues may be common in nature (Andreou et al., 2017; Puebla et al., 2007; Servedio, Doorn, et al., 2011).

However, modelling studies have cast doubt on magic traits’ efficacy at preserving assortative mating (Servedio and Bürger, 2014). Multiple studies have found that when two populations in secondary contact occupy different ecological niches, magic cues can only maintain species boundaries when there is a search cost (Boughman and Servedio, 2022; Yukilevich and Aoki, 2022). Search costs have also been shown to promote species coexistence without niche differentiation in a haploid model with spatial variation in carrying capacity (M’Gonigle et al., 2012), but both search costs and magic traits have received little attention from a tension zone perspective. This is because previous studies have used models that are haploid or lack spatial structure, precluding the existence of tension zones. Hence, here we simulate hybrid zones in continuous space, using populations of diploid individuals. Our goal is to determine whether there are conditions under which search costs or pleiotropy between mating cues, mating preferences, and/or hybrid viability traits can maintain stable tension zones on a homogeneous landscape between two populations occupying the same ecological niche.

To accomplish this goal, we develop an integrative framework for the modelling of secondary contact using continuous 2-dimensional space, individual-based interactions, and density dependence. In our simulations, starting populations can differ in loci that determine genetic incompatibilities, a mating display trait (the “cue”), and a mating preference (which determines the probability of accepting mates depending on their cues). Previous modelling studies investigating the effect of magic cues have used two island models where magic traits are incorporated by having one mating trait allele favoured on one island and another favoured on the other island (Boughman and Servedio, 2022; Servedio and Bürger, 2014; Yukilevich and Aoki, 2022). Here, we consider magic traits for which there is divergent viability selection on the mating traits. We include this in our simulations via pleiotropy between the mating trait loci and the loci encoding hybrid incompatibility. This falls under the framework used by Gavrilets (2004) and Kopp et al. (2018), but is distinct from scenarios in which divergent selection arises from adaptation to different niches in different parts of the species range. Magic cues in populations without niche differentiation can occur by exogenous selection—such as when hybrids are more susceptible to predation (Merrill et al., 2012; Noonan and Comeault, 2009), or by endogenous selection through Dobzhansky-Muller incompatibilities on mating trait loci (Qvarnström and Bailey, 2009).

A feature of our spatial model is that it allows for analysis of the phenotype distribution within a hybrid zone. Following Harrison and Bogdanowicz (1997), we categorize our simulation results as “bimodal” or “unimodal” based on the phenotype proportions at the centre of the hybrid zone. A hybrid zone is bimodal when most of the individuals at the centre possess a “parental” phenotype (one of the starting divergent phenotypes), and unimodal if they are mostly hybrids. Bimodality of a hybrid zone is considered an important indicator that the two populations function as distinct biological units since it shows their ability to remain distinct in sympatry (Jiggins and Mallet, 2000). Hence, bimodality has implications for species conservation (Fitzpatrick et al., 2008). Additionally, in bimodal hybrid zones, reinforcement is thought to be more likely since there are more interactions between the parental phenotypes (Coyne and Orr, 1997). Here, we simulate secondary contact with different parameter combinations specifying the fitness of hybrids, the strength of assortative mating, the genetic architectures of these traits, and the presence or absence of search costs. Each simulation outcome is categorized by the change in cline width over time and the bimodality of the hybrid zone at the end of the simulation. This allows us to determine which parameter combinations create stable species boundaries, and how changing any parameter affects the phenotype distribution within a hybrid zone.

## 2. Methods

Our model simulates secondary contact between two distinct populations. The model is based on the Hybrid Zone with Assortative Mating (HZAM) model (Irwin, 2020) and the sympatric version, HZAM-Sym (Irwin and Schluter, 2022). A key difference in the updated model, named HZAM-2D, is that the hybrid zone is now two-dimensional as opposed to a linear range in the HZAM model and no spatial component in the HZAM-Sym model. The current implementation is written in Julia (https://julialang.org; Bezanson et al. 2017) and is made available on GitHub (Irwin and Farley, 2025). HZAM-2D is an individual-based simulation of two starting populations (A and B) made up of diploid males and females (equal numbers of both populations and sexes to start the simulation). One or more loci—which follow rules of Mendelian inheritance and can take values of 0 or 1—determine the differences between the populations. Population A individuals start as 0/0 homozygous for all loci, and population B individuals start as 1/1 homozygous. Loci can be assigned to any of three traits: mating cue, mating preference, and hybrid incompatibility. Trait values are defined as the mean allelic value at loci for that trait.

Simulations take place on two continuous geographic dimensions (*x*, *y*) of length 1. At the beginning of each simulation, individuals are assigned random locations drawn from a uniform distribution, with population A restricted to the left half of the range (0 < *x* < 0.5), and population B restricted to the right half (0.5 < *x* < 1). The full range has a carrying capacity of *K* individuals (which includes both populations), and each population starts with 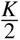 individuals. Carrying capacity also acts locally, such that females in low-density regions tend to have more offspring than those in high-density regions (see section 2.2 below). Each generation consists of a mating stage when females select male mates, a reproduction stage when the number of offspring is determined, and a survival to adulthood stage. Generations are non-overlapping, and individuals go through each stage once. The stages correspond to three different potential selective pressures: mate choice, density-dependent population regulation, and the reduced fitness of hybrids.

### 2.1. Mating

During the mating stage of the simulation, each female is presented with a single male at a time, with males sorted by geographic distance (the closest male presented first). The female can choose to accept or reject the presented male until she has found a mate or rejected every male within the cutoff distance (0.05). Acceptance probability is determined by the difference between the female’s mating preference trait value (*T_female_*) and the male’s mating cue value (*T_male_*). The probability of acceptance declines as their difference (|*T_female_* — *T_male_*|) increases, according to a Gaussian function with mean zero and standard deviation *σ_p_*. The standard deviation is determined by the strength of assortative mating (*S*_AM_):

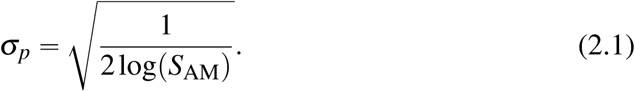

*S*_AM_ is defined as the ratio of the probability of a female accepting a male whose cue value matches her preference value to the probability of a female accepting a male whose mating cue value differs by 1 from her preference value. For example, when *S*_AM_ = 300, a female from population A is 300× more likely to accept a male from population A than a male from population B. When *S*_AM_ = 1 (no assortative mating), the standard deviation is taken to be infinite and when *S*_AM_ is infinite (i.e., perfect assortative mating, with no hybridization), the standard deviation is taken to be 10^−15^ (effectively zero).

### 2.2. Density-dependent population regulation

The number of offspring of each mated female is drawn from a Poisson distribution with a mean (*c*) adjusted to reflect local density dependence, a result of competition for resources. Local density (*N*_local_) at a female’s location (***u*** Ɛ [0, 1]^2^) is measured using unnormalized Gaussian kernel density estimation with standard deviation *σ_c_*:

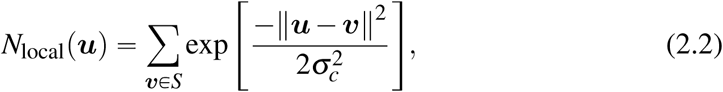

where *S* is the set of locations (in [0, 1]^2^) of all other individuals within 3*σ_c_* of ***u*** (for computational efficiency). The local carrying capacity at each location, *K*_local_, is calculated similarly, but assuming that *K* individuals are distributed evenly:

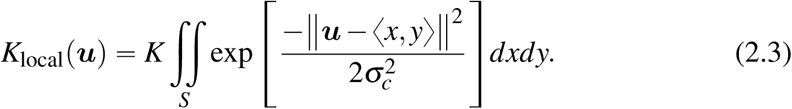

The expected mean number of offspring (*c*) for a female at location ***u*** was then calculated using the Beverton-Holt model—a discrete-time analogue for the logistic growth model (Beverton and Holt, 1993):

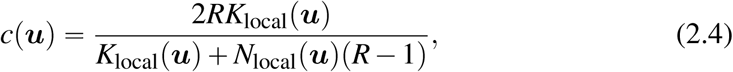

where *R* is the intrinsic growth rate of the population. In the simulations presented here, *R* = 1.1 (i.e. an expected 10% per generation growth rate when the population density is low) and *σ_c_*= 0.01. The sex of the offspring is assigned randomly.

### 2.3. Search cost

In a subset of simulations, there is an added reproductive cost *α* for each mate rejected, representing a cost of time or resources involved in interacting with the rejected individual and searching for another potential mate. With the added cost, (2.4) becomes

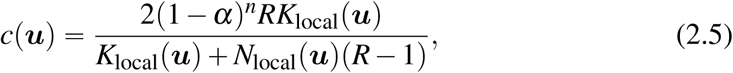

where *n* is the number of males that the female at location ***u*** rejected.

### 2.4. Survival probability

To model reduced hybrid fitness, hybrid individuals in the simulation can have a lower probability of survival to adulthood. Complete heterozygotes at all hybrid incompatibility loci (e.g. *F*_1_ hybrids) have survival probability *w*_hyb_, whereas pure homozygotes all survive to adulthood. The probability of survival is determined following the epistasis model of Barton and Gale (1993):

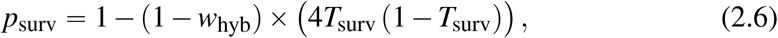

where *T*_surv_ is the hybrid incompatibility trait value—defined as the mean allelic value for the hybrid incompatibility loci. Note that while we refer to these as incompatibility loci, the cause of the low hybrid fitness can be ecological in nature, for instance if the loci encode a trait determining optimal resource characteristics and there is a valley in the resource availability distribution. Hence this is a model of divergent selection.

### 2.5. Dispersal

The dispersal distance of each surviving offspring is taken from a normal distribution with standard deviation *σ_d_* = 0.03. The dispersal direction is randomly assigned (i.e., an angle between 0 and 2*π*). If the dispersal vector added to the mother’s breeding location falls outside of the geographic range, the dispersal vector is reflected off the edge of the range.

### 2.6. Cline fitting and hybrid zone quantification

We use three measures to quantify simulation results: cline width, overlap, and bimodality. The mating cue is used as the basis for these calculations. We reason that such traits tend to be apparent to biologists and are used to assign individuals to species and to map the shapes of hybrid zones in empirical systems (whereas preferences and incompatibilities are more difficult to assess). Individuals with mating cue trait values (*T_cue_*) below 0.1 or above 0.9 are considered to possess a parental phenotype, and individuals with 0.1 ≤ *T_cue_*≤ 0.9 are deemed hybrids. Since all non-parental genotypes fall within the hybrid phenotype cutoff when there are fewer than four loci, this exact cutoff value only becomes relevant for our nine loci simulations.

Figure 1 shows how the cline width is calculated by fitting a sigmoidal curve to the average mating cue along five evenly-spaced horizontal transects. The average mating cue at each point along the transect is determined using Gaussian kernel density estimation with standard deviation *σ_c_* (see (2.2)). A sigmoidal curve is then fit to the average mating cue along each transect using the LsqFit Julia package. The cline width along a transect is defined here as in Barton and Hewitt (1985), as the inverse of the maximum gradient of the sigmoidal curve. The overall cline width is the mean of the cline widths along the five transects. Population overlap is defined here as the percentage of the range occupied by both parental mating cue phenotypes. This is calculated by measuring the population density (see (2.2)) for both populations at evenly spaced locations across the range in a 100 × 100 grid. The total population overlap is taken to be the proportion of sampled points where each of the two original mating cue phenotypes accounts for at least 10% of the population density.

**FIGURE 1.**
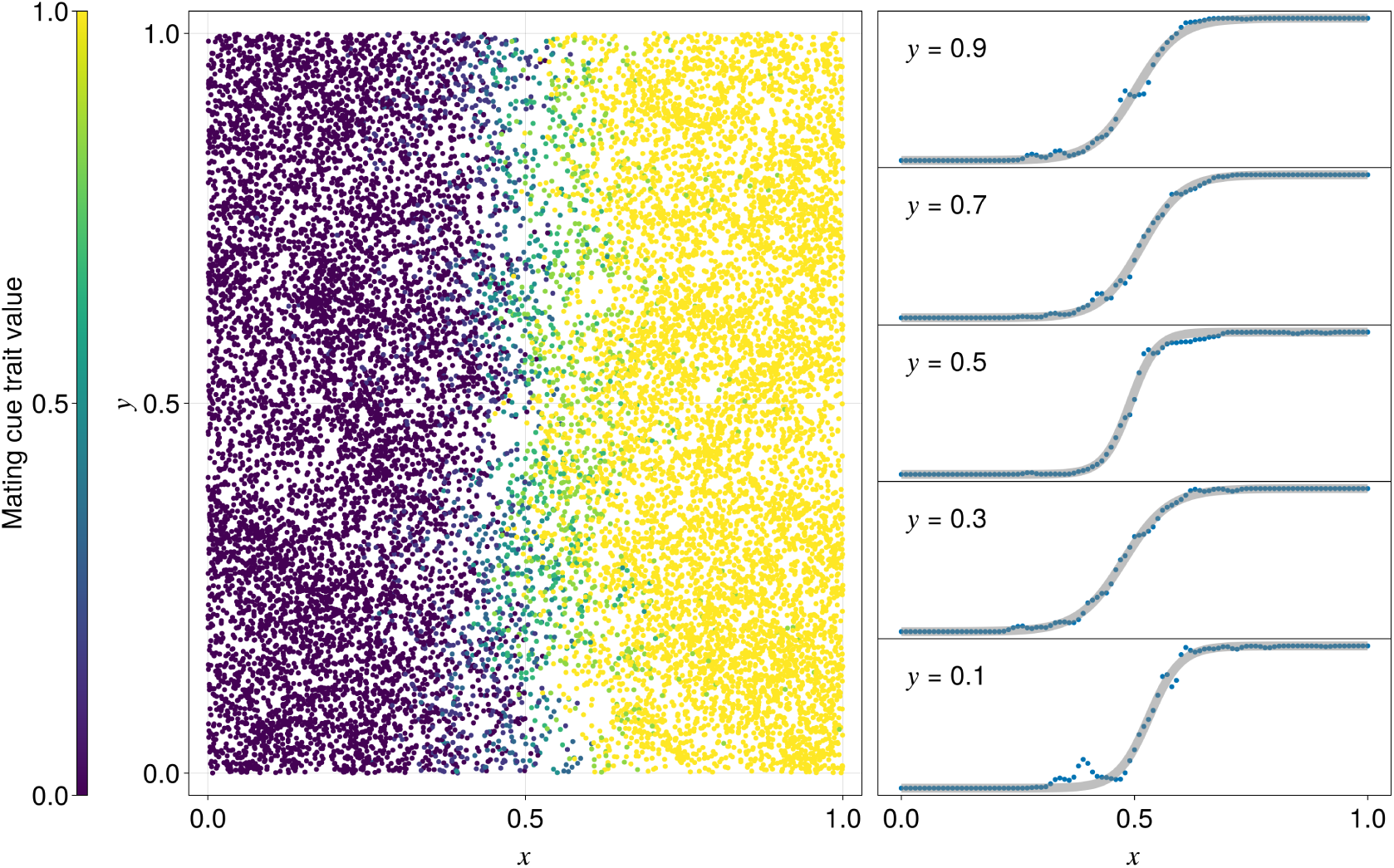
Cline width is calculated by fitting a sigmoidal curve to the average mating cue along five evenly spaced transects, then taking the mean width. The left panel shows the location and mating cue of every individual in one generation of a simulation, and the right panel shows the sigmoidal curves fit to the mating cue along each transect. The bimodality of this simulation was 0.11, meaning that at the centre of the hybrid zone, 11% of the population density was accounted for by parental phenotypes. The population overlap is 0.3%.

Unimodal hybrid zones are characterized by a gradual progression of intermediate phenotypes across the middle of the zone, whereas bimodal hybrid zones contain few intermediate phenotypes (Jiggins and Mallet, 2000). We quantify bimodality by calculating the population density (see (2.2)) of the parental mating cue phenotypes at the locations where the cline fit crosses 0.5 along each transect. The bimodality (*B*) at such a location is defined as the proportion of the local population density accounted for by parental genotypes, and the bimodality of the hybrid zone is taken to be the average across the five transects.

### 2.7. Running simulations and categorizing outcomes

Unless otherwise indicated, the simulations begin with 20,000 individuals and have three loci specifying each trait (cue, preference, and hybrid incompatibility), though these loci may overlap depending on the specified form of pleiotropy within each simulation. In all simulations, *R* = 1.1*, σ_c_*= 0.01, and *σ_d_* = 0.03, ensuring an equilibrium population size between 16,000 and 20,000 individuals depending on *w*_hyb_, *S*_AM_ and *α*. These values were chosen so simulations could run efficiently while ensuring that the population was large enough such that random chance played only a small role in determining the simulation outcomes. The mean dispersal distance was selected to be as small as possible (to make measurements of cline width and population overlap more precise) while still ensuring a roughly even distribution of individuals across the landscape. Most simulations ran for 1,500 generations, with the cline width measured every 50 generations.

Simulation sets were run using all combinations of *S*_AM_ Ɛ {1, 3, 10, 30, 100, 300, 1000, complete} and *w*_hyb_ Ɛ {0, 0.1, 0.2, 0.3, 0.4, 0.5, 0.6, 0.7, 0.8, 0.9, 0.95, 0.98, 1}. We used a four-step process to categorize the simulation outcomes. (1) Simulations were determined to be *extinctions* (of one starting population) if one parental mating cue phenotype was absent and the other accounted for over half of the individuals at the end of the simulation. (2) The simulations that were not extinctions were categorized as *blended* if the cline width was increasing over the last 1,000 years of the simulation or if the cline width ever exceeded the width of the simulated range. To determine if the cline width was increasing, we performed a Mann-Kendall test on the cline width measurements (recorded every 50 generations) with a significance level of 5%. (3) Each simulation with a stable cline width was classified into an outcome type determined by the hybrid zone bimodality (*B*) at the end of the simulation. The three basic outcome types were (A) unimodal hybrid zone (*B* < 0.5), (B) bimodal hybrid zone (0.5 ≤ *B* ≤ 0.95), and (C) overlap zone (*B* > 0.95). (4) Finally, overlap zones were further divided into “narrow overlap zones” (overlap < 10*σ_d_*) and “broad overlap zones” (overlap ≥ 10*σ_d_*).

## 3. Results

The simulations show that the persistence of distinct mating types after secondary contact is highly dependent on the genetic architecture of the mating trait(s), in addition to the strength of assortative mating and hybrid fitness. Including search costs and/or pleiotropy for combinations of mating cue, mating preference, and hybrid incompatibility traits can in many cases prevent blending, and there are qualitative differences in outcome between various combinations of these factors. To demonstrate these differences, we start by comparing four scenarios: (1) cue, preference, and hybrid incompatibility are all encoded by separate loci (no pleiotropy); (2) the same, except there is a mate search cost such that female fecundity is reduced 5% for each male rejected as a mate (no pleiotropy with search cost); (3) cue and hybrid incompatibility are encoded by the same loci, but preference is separate (magic cue); and (4) the same, but with a 5% mate search cost (magic cue with search cost). These four example simulations were all run with 90% hybrid genetic compatibility (*w*_hyb_ = 0.9) and a 10× preference of females for males of their own population compared to males of the other population (*S*_AM_ = 10).

In the no pleiotropy simulation under these conditions (Fig. 2A), the association between preferences and cues rapidly breaks down over the first few hundred generations, allowing intermediate mating phenotypes to spread from the centre of the contact zone. The result is a preference distribution that is genetically unimodal and spatially uniform (Fig. 2A, top panels). After 2,000 generations, intermediate preferences have caused all the cue loci to become fixed, resulting in an intermediate cue phenotype (Fig. 2A, bottom panels). At that point, variation in preference persists, as it is no longer under selection. The collapse in assortative mating contrasts with the persisting stable cline in the incompatibility loci (which causes the low-density region in the rightmost panels of Fig. 2A). This simulation illustrates how without search costs or pleiotropy, the boundary between parental mating types cannot be maintained at these moderate levels of assortative mating and hybrid fitness (as found in previous work).

**FIGURE 2.**
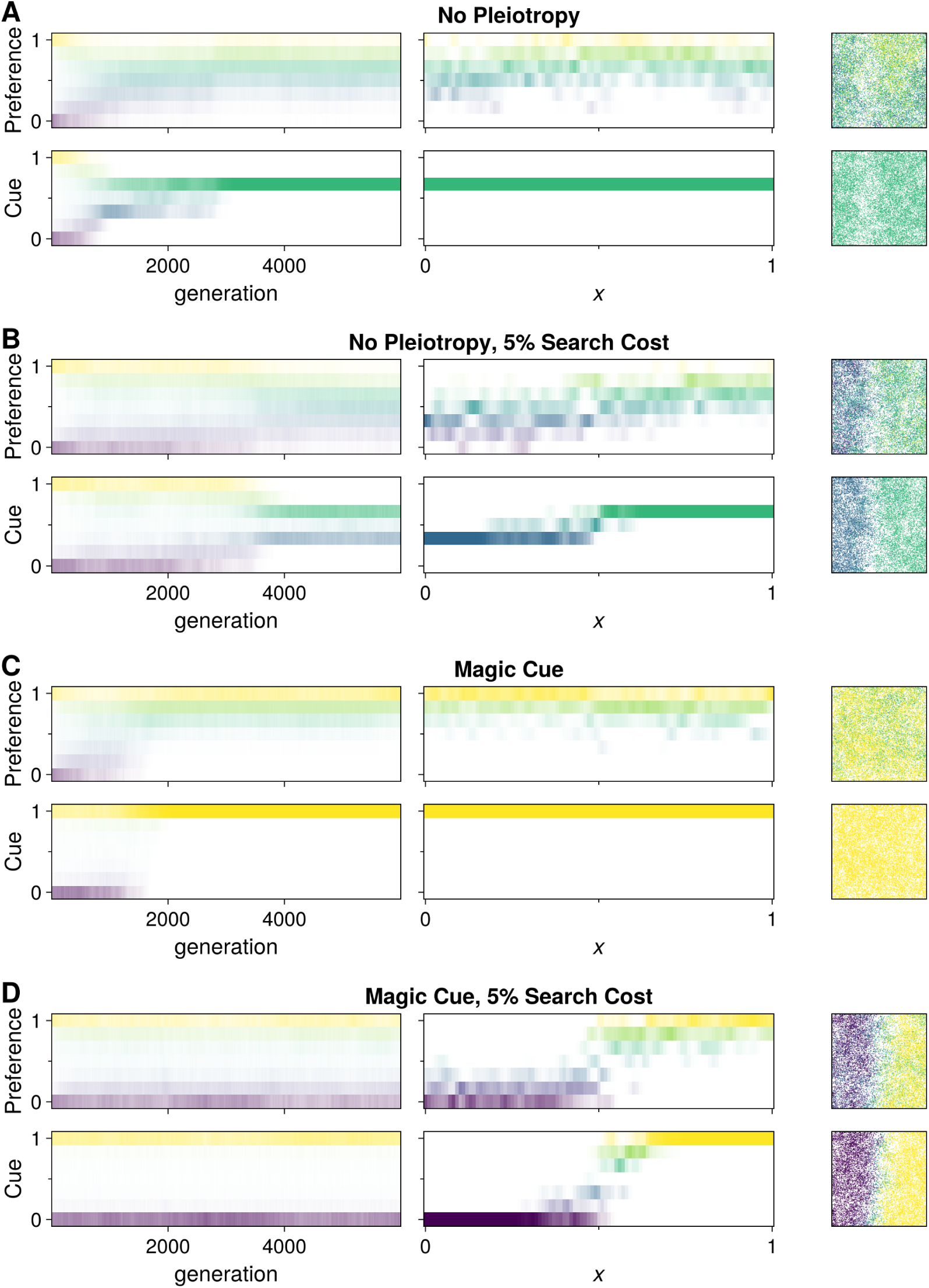
Four example simulations illustrating possible outcomes. All simulations were run with *S*_AM_ = 10, and *w*_hyb_ = 0.9. For each simulation, mating preference and mating cue are shown as phenotype frequencies changing over time (left panels), phenotype densities along the *y* = 0.5 transect after 6,000 generations (middle panels), and spatial distributions of individuals after 6,000 generations (right panels).

When we start with the same conditions as above but add a search cost of 5% fitness per rejected potential mate (Fig. 2B), the outcome switches to a hybrid zone that is relatively stable for many generations, although there is a reduction in differentiation. Initially, the preferences gradually approach intermediacy (Fig. 2B top panels), but at a much slower rate than in the no pleiotropy simulation. Weak selection on mating preference facilitates gene flow, ensuring that preferences at any given location are unimodal. This causes strong selection on the mating cue to align with the local average preference and makes cue polymorphism unstable. The mating cue cline collapses into a single-locus cline, but it cannot collapse further without creating areas with overlapping phenotypes. The search cost causes selection on mating preferences to align with local cues, preventing the preference cline from collapsing any further than the cue cline. This results in stable mating cue and preference distributions at the end of the simulation.

In the magic cue simulation (Fig. 2C), in which loci encoding the cue also encode low hybrid fitness, initial rare hybridization and gene flow eventually cause collapse of the mating preference cline, causing more buildup of hybrids with intermediate and low-fitness cues. The fitness valley causes divergent selection, leading to the extinction of one of the mating cue types (which one is arbitrary and varies between simulation runs). In watching the simulations, we observe that the cline in preferences expands over time, which causes the mating cue cline to slowly widen. Because mating cue and hybrid fitness have a shared genetic basis, individuals with intermediate cues have low fitness, resulting in an expanding “population sink”. Once the hybrid zone grows too close to an edge of the simulated range, the rate of gene flow becomes insufficient to maintain the position of the hybrid zone. The shift of the hybrid zone toward the smaller population drives the extinction of one of the mating types, and the males with the other parental mating cue phenotype quickly outcompete the low-fitness hybrids.

Finally, in the simulation with both the magic cue and the search cost (Fig. 2D), we see a broad stable cline in the mating preference and a narrow stable cline in the mating cue. Unlike in the previous simulation, pleiotropy with the incompatibility loci prevents the buildup of mating cue phenotypes with intermediate values. The search cost constricts the preference cline, since there is selection against females with preferences misaligned with the local cues. The result is a stable, well-defined mating-type boundary that would not exist without both the magic cue and the search cost.

In Figure 3, we summarize how search costs and different pleiotropic combinations of traits interact with hybrid incompatibility and assortative mating to determine the outcome of hybridization between two populations meeting in secondary contact. We ran simulations with 936 different parameter combinations, and show for each parameter combination the first outcome type to occur three times (see Methods for criteria). In the no pleiotropy simulations (Fig. 3A), we see that about half of the studied parameter combinations of assortative mating strength (*S*_AM_) and hybrid fitness (*w*_hyb_) result in blending of the two starting populations. Broad overlap zones occur only with extreme values of assortative mating strength (i.e., infinite when hybrid fitness is greater than 0.6, and 300× or higher at lower values of hybrid fitness). Hybrid zones are only maintained when hybrid fitness is low, e.g. 0.6 or lower when *S*_AM_ ≤ 300. The boundary between a blended outcome and one of the outcomes in which divergence is maintained angles from lower left to upper right on this graph, from about *S*_AM_ = 3 and *w*_hyb_ = 0.3 to *S*_AM_ = 1000 and *w*_hyb_ = 0.7. It is likely that many natural hybrid zones do not have such extreme combinations of low hybrid fitness and strong assortative mating, raising the question of what other factors are involved. We now turn to how search costs and pleiotropy affect these outcomes.

**FIGURE 3.**
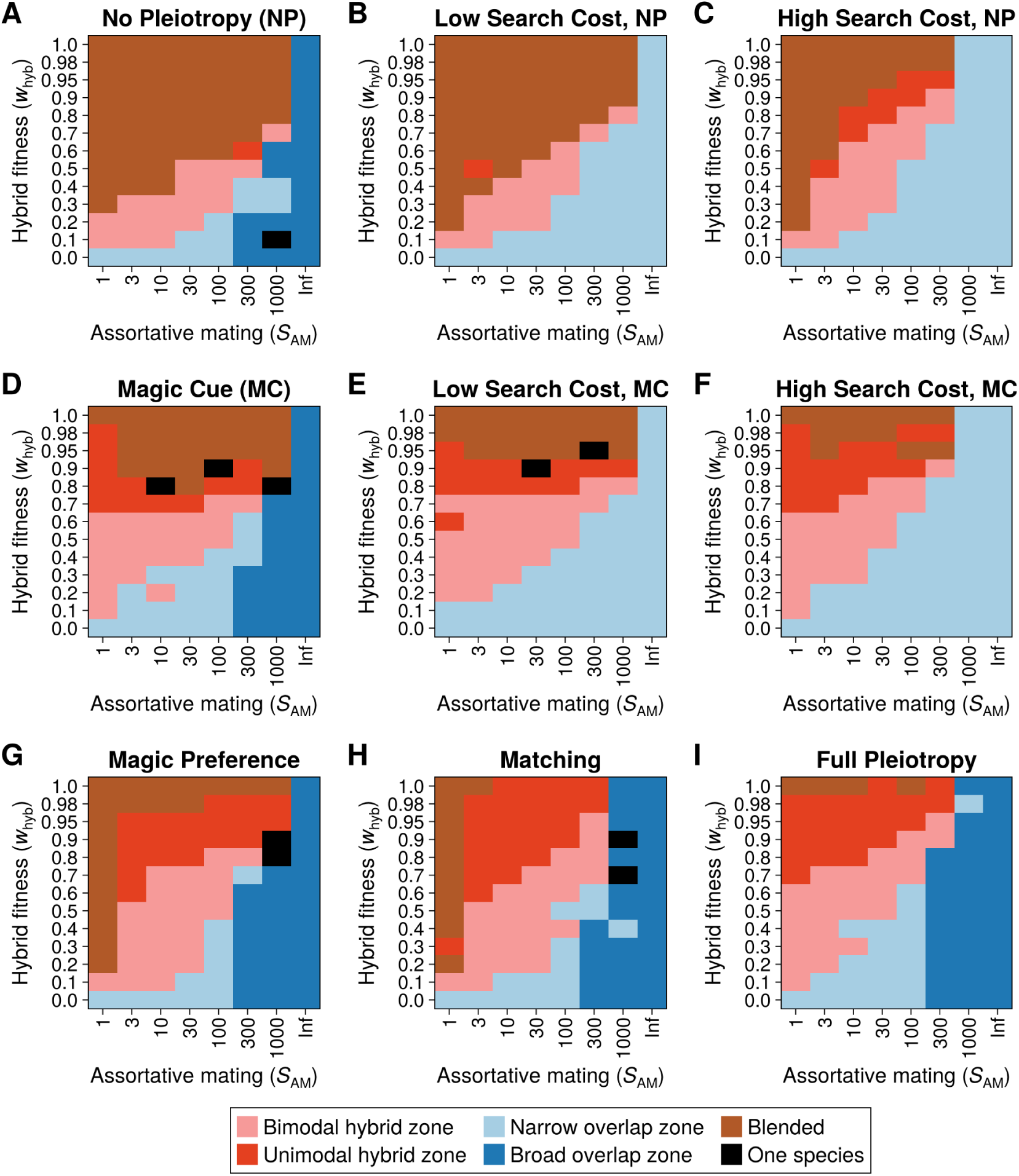
Simulation outcomes (after 1500 generations) as determined by the distribution of mating cue phenotypes, under different combinations of strength of mating preference (*S*_AM_) and hybrid genetic incompatibility (*w*_hyb_), and nine different scenarios regarding genetic architecture and search costs. For “full pleiotropy,” all three traits (incompatibilities, mating cue, and mating preference) are determined by the same 3 loci, whereas for “no pleiotropy” these three traits are each determined by different sets of 3 loci. For “magic cues/preferences,” the incompatibility and cue/preference traits are determined by the same loci, respectively, and for “matching” the cue and preference are determined by the same loci. Low and high search costs are 1% and 5% fitness reductions per rejected male, respectively.

When we add search costs to the no pleiotropy simulations (Fig. 3B,C), the boundary between the blended outcome and the hybrid zone outcome shifts upward and to the left, toward higher values of hybrid fitness and lower values of assortative mating strength. This is for two reasons: (1) when hybrids are rare, they pay a large search cost because they reject many potential mates, reducing the fitness of hybrids and thereby inhibiting their buildup, and (2) when either species is rare, for instance when they’ve dispersed across a hybrid zone, they reject many potential mates and therefore pay a large search cost, limiting the ability of a rare species to persist locally. These effects are modest at low search costs (Fig. 3B), and much more apparent at high search costs (Fig. 3C). In the latter case, combinations such as *S*_AM_ = 10 and *w*_hyb_ = 0.8, a modest level of reproductive isolation, now result in a maintained hybrid zone. We also note that the “broad overlap” outcome is not seen in the search cost simulations, because the cost of rejecting mates at high levels of assortative mating impacts the locally rarer species, preventing sympatric coexistence. We anticipate that in real species interactions, negative interactions between species likely go down at high strengths of assortative mating, lowering the impact of search costs and allowing more broad coexistence than predicted by our models of constant search cost. Search costs had a much larger effect in the one-locus simulations, where almost all parameter combinations resulted in stable clines even when the search cost was only 1% (see Fig. A-1). This echoes what is observed in Figure 2C, where the search cost leads to a three-locus cline collapsing down to a stable single-locus cline.

We now turn to the effect of pleiotropy on these outcomes. Magic cues (pleiotropy between mating cues and incompatibility loci; Fig. 3D) shift the boundary between blended outcomes and stable hybrid zones upward along the hybrid fitness axis, in comparison with the no pleiotropy case. Now, the boundary between a blended outcome and a stable hybrid zone outcome occurs at about *w*_hyb_ = 0.8, much higher than in the no pleiotropy case (Fig. 3A), especially at low values of *S*_AM_. When hybrid genetic incompatibility is lower than this value, the divergent viability selection on the mating cue can counteract the pull of the collapsing preference cline, preventing the mating cue cline from collapsing within 1,500 generations. These results were similar across different numbers of loci (Figs. A-1 and A-2). Adding a mate search cost (Fig. 3E–F) reduces the number of blended outcomes, with a larger search cost having a bigger impact in this way.

While we have seen above that magic cues (Fig. 3D–F can stabilize hybrid zones compared to the no pleiotropy case (Fig. 3A-C), magic preferences (pleiotropy between mating preferences and incompatibility loci) are even more effective (Fig. 3G). We now see that hybrid zones are maintained for quite modest levels of reproductive isolation, for instance when *w*_hyb_ = 0.95 and *S*_AM_ = 3. Blending under these conditions is prevented because the pleiotropy means there is direct selection against intermediate preferences, maintaining a preference cline, and the divergent preferences place divergent selection on the mating cue, maintaining the cline in cue as well. We note that Fig. 3G shows a blending result along the left column because *S*_AM_ = 1 means there is no preference among potential mates, allowing the cue-encoding loci to flow relatively freely and resulting in blending of the cue. This is not seen in the left (*S*_AM_ = 1) column of the magic cue results (Fig. 3D) because in that case the cue is pleiotropic with incompatibility loci, such that low hybrid fitness maintains the cue cline even without mate choice.

In addition to examining scenarios where preferences and cues are encoded by different loci, we can look at what happens when they are pleiotropic—such that the mating preference and cue can be considered a single trait. Following the terminology from Kopp et al. (2018), we call the latter case “assortative mating by matching”. Fig. 3H shows the results for such a case of cue-preference pleiotropy when incompatibilities are encoded by other loci, and Fig. 3I shows full pleiotropy between the preference, cue, and incompatibility loci. Results in these two cases are nearly identical, except for two differences. The first is that when the hybrid incompatibility trait is separate and *S*_AM_ = 1, the “assortative mating by matching” setup is identical to the leftmost columns of the “magic preference” (Fig. 3G) and “no pleiotropy” (Fig. 3A–C) setups. The second is that broad overlap zones occur more readily under full pleiotropy. This is because there are more hybrids produced when hybrid incompatibility is separated from the mating traits since not all hybrids possess genetic incompatibilities. A similar pattern is seen in how there are fewer broad overlap zones in the magic preference simulations (Fig. 3G) when hybrid fitness is high (0.95 ≤ *w*_hyb_ ≤ 0.98). More hybrids (which are categorized as such by their mating cues) are produced since a female’s mating preference may not align with her mating cue loci. This results in a unimodal hybrid zone dominated by intermediate phenotypes under the same assortative mating and hybrid fitness conditions that created broad overlap zones in the matching and full pleiotropy simulation sets. So not only does the genetic architecture impact the overall stability of mating trait differentiation, but it also shapes the phenotype distribution within stable hybrid zones.

We categorized extinction outcomes as the complete absence of one of the parental mating cue phenotypes and over 50% of the population consisting of the other mating cue phenotype. Extinctions occurred at a low frequency under a wide range of parameter combinations, but were much more likely when assortative mating was strong. Among the first 3 simulation runs on all parameter combinations, 48% of the extinction events occurred with 1000× assortative mating. This is compared with 1% for no interbreeding and 18% with 300× assortative mating. No extinction events occurred when there was 1× assortative mating. The reason for this is that strong assortative mating leads to broad regions of population overlap. Population overlap between two populations occupying the same niche is unstable since the rarer population suffers from interbreeding with the more numerous population. This leads to rapidly shifting population distributions and makes extinction much more likely.

Although Figure 3 shows the magic cue as being fairly effective at maintaining population boundaries, Figure 2C demonstrates that the clines may not be as stable as they appear. In Figure 2C, there was an apparently stable mating cue cline for the first 1,500 generations (the timescale used in Figure 3) that rapidly collapsed at the 2,000 generation mark. To further investigate this, we re-ran the magic cue simulations and categorized the results based on the mating preference rather than the cue. Ordinarily, the preference and cue clines align closely. But pleiotropy between the mating cue and the incompatibility loci means that the cue cline width can remain steady even while the preference cline gradually collapses. This can create the illusion of a stable boundary, but once the preference cline grows wide enough, the cue cline will collapse as seen in Figure 2C. The results in Figure 4 show that preference clines under magic cues are as unstable as the cue clines under no pleiotropy in Figure 3. This indicates that the combination of a magic cue and a search cost is only slightly more effective at maintaining stable population boundaries long-term than search costs alone.

**FIGURE 4.**
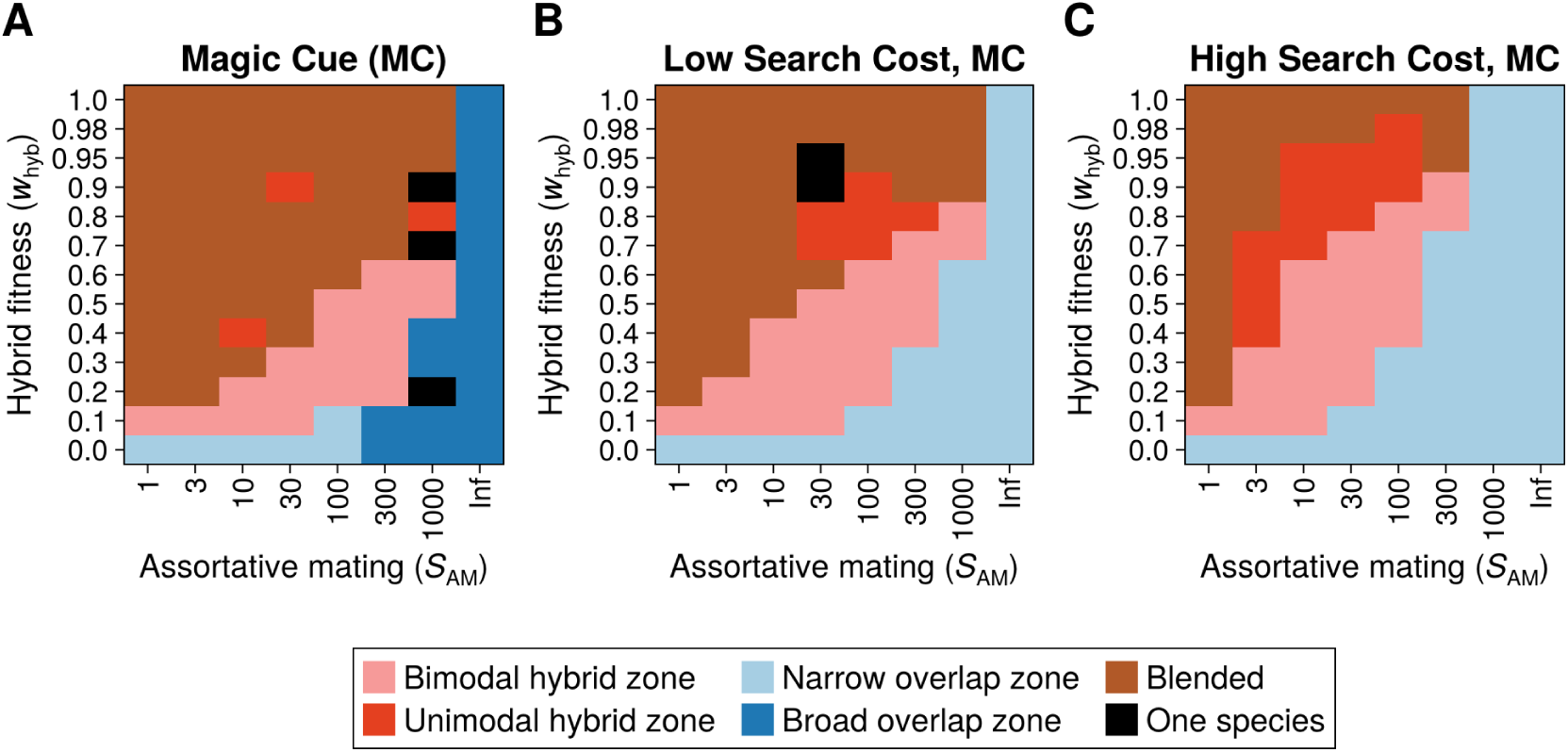
Magic cue simulation outcomes (after 1500 generations) as determined by the distribution of mating preference phenotypes, under different combinations of strength of mating preference (*S*_AM_) and hybrid genetic compatibility (*w*_hyb_), and three different search cost strengths (0%, 1%, and 5% fitness reduction per rejected male).

## 4. Discussion

Hybrid zones arising from secondary contact are common in nature, and some are thought to have existed for thousands of years (Gwee et al., 2025; Moore and Buchanan, 1985). In some of these ancient hybrid zones, assortative mating is still present (Haas et al., 2010; Haavie et al., 2004; Wiebe, 2000). This leads to the question of how assortative mating is sustained in the presence of gene flow between species. Many hybrid zones are narrow clines maintained by a balance of dispersal of parental genes into the hybrid zone and selection against hybrid genotypes within the hybrid zone; these are called “tension zones” (Barton and Hewitt, 1985). The spatial nature of a tension zone allows for the formation of a “hybrid sink”—an area of low population density impeding gene flow (Barton, 1986), or a “hybrid bridge”—a series of intermediate phenotypes linking two populations with otherwise low direct mating compatibility (Broyles, 2002). The spatial structure is particularly important when looking at assortative mating, as the fitness of a mating trait phenotype will vary across the landscape according to local mating preference/cue distributions (Bridle et al., 2006). Single-locus clines can be stabilized by genetic incompatibility or selection against rare types (Gavrilets, 1997). But when mating cues and preferences are encoded by separate loci, mating trait clines are prone to collapse due to the weak selection on preference genes.

Previous models of secondary contact with separate mating preference and cue traits used haploid or two-island models to examine the effects of search costs and magic cues (Boughman and Servedio, 2022; M’Gonigle et al., 2012; Servedio and Bürger, 2014; Yukilevich and Aoki, 2022). These studies found that magic cues on their own do little to maintain species boundaries, but can promote coexistence when combined with search costs and local adaptation. However, due to the nature of haploid and two-island models, there is a gap in the literature as to whether these factors allow for stable tension zones to form. Here, we have examined a model of secondary contact on a two-dimensional homogeneous landscape that can include magic traits and search costs, allowing inference of the effect of each factor and their combination on the stability and bimodality of mating cue distributions in contact zones.

Results indicate that magic cues can create temporarily stable mating cue clines, but fail to stabilize preference clines—ultimately leading to the loss of assortative mating. Search costs result in stable clines in a wide set of circumstances when there is a small number of loci, but otherwise require very strong assortative mating. The combination of magic cues and search costs is more effective than magic cues, but is not much better than search costs without pleiotropy at maintaining stable mating preference clines (Fig. 4). Search costs do not require pleiotropy to maintain stable tension zones, and can do so even without hybrid genetic incompatibility when cues or preferences are encoded by a single locus (Fig. A-1B,C). We find that small changes in assortative mating strength can change the phenotype distribution of a contact zone from a unimodal hybrid zone with predominantly hybrids to an extensive region of population overlap with few hybrids. The assortative mating threshold needed to create population overlap is dependent on the genetic architecture of the mating traits, and is typically on the order of 1000×.

## Magic cues

One difficulty in magic trait research is that its definition is understood differently in the contexts of speciation and hybrid zones. Magic trait speciation models come in two main varieties: (1) two-island models with local adaptation (Thibert-Plante and Gavrilets, 2013), and (2) sympatric models with resources distributed along a trait axis (Dieckmann and Doebeli, 1999; Verzijden et al., 2005). Both of these build off of earlier speciation models, such as the one developed by Maynard Smith (1966), that do not include assortative mating. Although most of these models use full pleiotropy (Dieckmann and Doebeli, 1999), some examine magic cues as well (van Doorn et al., 2004). Magic preference speciation models are less common, but they have been used in the context of sensory drive (Kawata et al., 2007).

In hybrid zones, another natural way to model divergent selection on mating traits is hybrid inviability (Dobzhansky, 1937; Muller, 1942). Selection against hybrids causing postzygotic reproductive isolation forms the foundation of much of hybrid zone theory and does not require local adaptation or resource partitioning (Barton, 1979; Barton and Gale, 1993). This has led researchers working on hybrid zones to adopt a broader definition of magic traits that includes mating trait loci involved in intrinsic hybrid incompatibilities (Kopp et al., 2018). Proving the existence of this type of magic trait will generally be more difficult than a niche-based magic trait since there may not be any external indicators of low fitness. Nevertheless, sexual selection may cause loci encoding mating traits to diverge more rapidly than other parts of the genome in allopatry, making them more prone to intrinsic incompatibility upon secondary contact (Sloan et al., 2023). Furthermore, a model for intrinsic hybrid incompatibilities can be extended to other selection pressures against hybrids, such as from predators. This is particularly relevant to hybrid zones between parental populations that have evolved to mimic different species, as the hybrid phenotype will break the mimicry and suffer from increased predation (Jiggins, Naisbit, et al., 2001; Symula et al., 2001).

Given the emphasis on magic cues (cue-fitness pleiotropy) in the speciation literature, one notable result from our study is that other types of pleiotropy tend to be more effective at preventing blending over much of the parameter space. In particular, cue-preference pleiotropy is more effective than magic cues at moderate levels of assortative mating strength and high levels of hybrid fitness. Extending this to full pleiotropy (by adding hybrid fitness) changes the results relatively little. Additionally, magic preferences produce similar results as full pleiotropy since the preference exerts direct selection on the mating cue (see Fig. 3). Since the effects of magic preferences are easily explained, previous modelling studies have, as we have here, focused on magic cues (Servedio and Bürger, 2014; Yukilevich and Aoki, 2022). Our results show that hybrid inviability magic cues can create temporarily stable mating cue clines, provided there is a moderate (> 10%) reduction in hybrid genetic compatibility. But crucially, magic cues do not stabilize mating preference clines since there is no selection on preference outside of the contact zone. An expanding preference cline will create selection for intermediate mating cues, which reduces the fitness of the population and eventually leads to the extinction of one mating cue type. This is a different outcome from what occurs in a local adaptation magic trait model. In that context, magic cues can maintain some mating cue divergence (Servedio and Bürger, 2014; Yukilevich and Aoki, 2022). This is because, unlike hybrid inviability, local adaptation creates a balance between the two populations and prevents either population from replacing the other. These differences underscore the importance of not only the strength but also the type of divergent selection in determining the outcome of secondary contact with magic traits.

## Search costs

Mate search costs occur when choosier females run the risk of having fewer offspring due to depleted energy stores, increased risk of predation, or bearing young at a suboptimal time (Alatalo et al., 1988; Forsgren, 1992; Watson et al., 1998). Search costs are known to suppress sympatric speciation (Gavrilets, 2004; Otto et al., 2008), and can also help maintain species boundaries after secondary contact (Boughman and Servedio, 2022; M’Gonigle et al., 2012). Our results show that search costs can effectively maintain stable tension zones under a wide range of conditions when one or both mating traits are encoded by a single locus (see Fig. A-1). When cue is encoded by a single locus and preference is encoded by multiple loci, weak selection on mating preference creates broad clines with little overlap in preference phenotypes. As demonstrated by Lande (1982), this ensures that at each location there is a single locally optimal cue phenotype. Selection toward a local optimum prevents mating cue polymorphism and, in doing so, stops single-locus cue clines from expanding. Search costs create selection on mating preference within and away from the contact zone, resulting in stable clines for both cue and preference (see Fig. 2B and Fig. A-3B). When multiple loci encode a mating cue trait, what can happen is that most of the loci become fixed, leaving a single-locus cue cline and a broad multilocus preference cline (see Fig. 2B). This shows how a hybrid zone arising from secondary contact can settle into a stable equilibrium where there is a sharp change in mating cue on a single locus, and an imperceptible gradient in mating preference. A similar pattern is observed when preference is encoded by a single locus (see Fig. A-3A), because alleles for a single-locus preference are less likely to introgress since the effective per-locus selection created by search costs is much higher than in the three-loci case. These results generalize to scenarios where the mating cue is encoded by multiple non-additive loci (see Fig. A-3C).

Search costs are likely present whenever there is assortative mating, but quantifying the costs is difficult. Across diverse taxa, experimental studies have shown that search costs influence mate choice (Alatalo et al., 1988; Booksmythe et al., 2008; Forsgren, 1992). Our results demonstrate that even a search cost as low as 1% per rejection of a potential mate can have an appreciable effect on cline stability, but it remains unknown if this value is realistic. M’Gonigle et al., 2012 found that search costs can create stable species boundaries in a haploid model when there is spatial variation in the resource carrying capacity. Here, we’ve shown in a diploid model that search costs can result in a stable tension zone on a uniform landscape. Previous studies using two-island models have found that search costs on their own fail to maintain species boundaries, but are effective in combination with a magic trait tied to local adaptation (Boughman and Servedio, 2022; Yukilevich and Aoki, 2022). However, the lack of spatial structure may in some cases exaggerate the effects of search costs. In a spatially explicit model, hybrids are concentrated in the middle of the contact zone where they are less affected by search costs. This can lead to a “hybrid bridge” effect, reducing the impact of search costs. Although search costs can result in stable tension zones in our spatial model, this requires both reduced hybrid fitness and strong assortative mating when traits are encoded by multiple loci. We found that the combination of search costs and magic cues was only slightly more effective than search costs without pleiotropy.

## Phenotype distribution

As well as varying in cline width, hybrid zones in nature differ dramatically in their phenotype distributions. Harrison and Bogdanowicz (1997) categorized hybrid zones based on their bimodality, where unimodal hybrid zones consist primarily of hybrids at their centre, and bimodal hybrid zones consist primarily of parental phenotypes. It is common for bimodality to vary even along a single hybrid zone that extends across a landscape, with mostly parental phenotypes along some transects and mostly hybrids along others (Howard, 1986; Szymura and Barton, 1986). We found that when traits are encoded by multiple loci, a small change in assortative mating strength can create an abrupt transition from a broad unimodal hybrid zone to an extensive overlap zone with few hybrids. This threshold is typically on the order of 1000× assortative mating strength, and is dependent on the genetic architecture of the mating traits. For a discussion of the conditions necessary for broad population overlap, see Irwin (2020) and Irwin and Schluter (2022). In nature, assortative mating strength is often determined by environmental conditions. Changes in the sensory environment will affect individuals’ abilities to differentiate mates (Seehausen et al., 2008), and resource abundance and predation risk determine the time available for searching for a mate (Booksmythe et al., 2008; Byers et al., 2006). Based on our results, even a small environmental effect on assortative mating strength could cause a hybrid zone to shift to dramatically different phenotype distributions.

When assortative mating strength is fixed (as it is in our model), hybrid zones are often unstable without magic traits, search costs, or preference-cue pleiotropy. This, however, may not hold when choosiness is an evolvable trait. Reinforcement is the process by which low hybrid fitness causes the evolution of increased prezygotic reproductive isolation (Butlin, 1987; Servedio and Noor, 2003), and it’s thought to occur more readily in hybrid zones with population overlap, since the two populations interact directly rather than through hybrid intermediaries (Cain et al., 1999; Jiggins and Mallet, 2000). Our results suggest that population overlap requires either preference-cue pleiotropy or substantial pre- and post-zygotic reproductive isolation. This would suggest that the parental phenotype distribution within contact zones is unfavourable to reinforcement under most conditions, but further research is needed.

## Conclusions

Across diverse taxa, hybrid zones often involve traits that appear to act as mating signals, such as song or coloration (DeRaad et al., 2023; Nürnberger et al., 2005). But often, little to no assortative mating is detected within the contact zone (Brelsford and Irwin, 2009; Moore, 1987). To further confuse matters, in some of these hybrid zones, there is assortative mating (as determined by mate-choice experiments) outside of the contact zone (Pascal et al., 2023; Ritchie et al., 1989). So why does assortative mating break down completely within contact zones, and what maintains mating trait differences outside of them? A key obstacle here is that the strength of assortative mating is difficult to measure in hybrid zones. Identifying genes encoding preferences is typically not possible, so studies rely on linkage between mate preferences and either phenotypic traits or the genes thought to be responsible for them (Haas et al., 2010; Nürnberger et al., 2005). Hence, assortative mating is easier to detect when preferences and cues are pleiotropic. This occurs in some hybrid zones where the mechanism for assortative mating is phenotype-matching (Conte and Schluter, 2013; Johnson, 1982; Kronforst et al., 2006), but it is far from universal.

Unlike in the pleiotropic case, separate preference and cue clines cannot be maintained by assortative mating alone (Felsenstein, 1981; Kirkpatrick and Barton, 1997; Kopp et al., 2018). In this paper, we’ve explored a few mechanisms that can help stabilize mating trait clines: magic cues, magic preferences, and search costs. Magic cues can maintain temporarily stable mating cue clines, but fail to stabilize preference clines. This ultimately leads to a collapse in the cue cline, but the process can take hundreds, if not thousands, of generations. Magic preferences provide a compelling way for both cue and preference clines to remain stable under preference/cue assortative mating. But although the theory behind them is well understood, it is far from clear how common they are in nature (Maan and Seehausen, 2012). The third mechanism we’ve discussed here is search costs. In our model, search costs maintain mating trait clines under a broad range of conditions unless the mating cue is encoded by multiple additive loci (in which case, low hybrid fitness and strong assortative mating are needed as well).

We’ve shown here how search costs can result in a narrow single-locus mating cue cline and a very broad multilocus preference cline. In such a case, the variation in mating preference would obscure assortative mating within the contact zone. Thus, a tension zone can be maintained by sexual selection alone, even when assortative mating is not detectable within the contact zone. Unlike magic traits, search costs require very weak assumptions. One only needs to assume that energy is expended in rejecting mates and that expended energy translates to lower reproductive output (Real, 1990). The generality of search costs makes them a credible mechanism for preserving assortative mating in a hybrid zone, but there is still much we don’t know. Here we’ve shown how assortative mating can be sustained after secondary contact with and without magic traits or preference-cue pleiotropy. Our results demonstrate the importance of not only the strength but also the type of selection acting on mating traits and highlight the need for more data on mating preferences and search costs.

## Author contributions

**Jack Farley:** conceptualization (equal), data curation (lead), formal analysis (lead), investigation (equal), methodology (equal), software (equal), visualization (lead), writing – original draft (lead), writing – review and editing (equal). **Darren Irwin:** conceptualization (equal), formal analysis (supporting), funding acquisition (lead), formal analysis (supporting), investigation (equal), methodology (equal), project administration (lead), resources (lead), software (equal), supervision (lead), validation (lead), visualization (supporting), writing – original draft (supporting), writing – review and editing (equal).

## Acknowledgements

We thank Jessica Irwin and members of the Irwin lab for discussion. This research was supported by a Discovery Grant (RGPIN-2023-04300) from the Natural Sciences and Engineering Research Council of Canada.

## Conflict of interest statement

The authors declare no conflicts of interest.

## Data availability statement

The software used for the simulations is available in https://github.com/jrfarley/HZAM-2D/releases/tag/v2.0.1-review. The data underlying this article will be archived in the Dryad digital repository before publication.

## Appendix

**FIGURE A-1.**
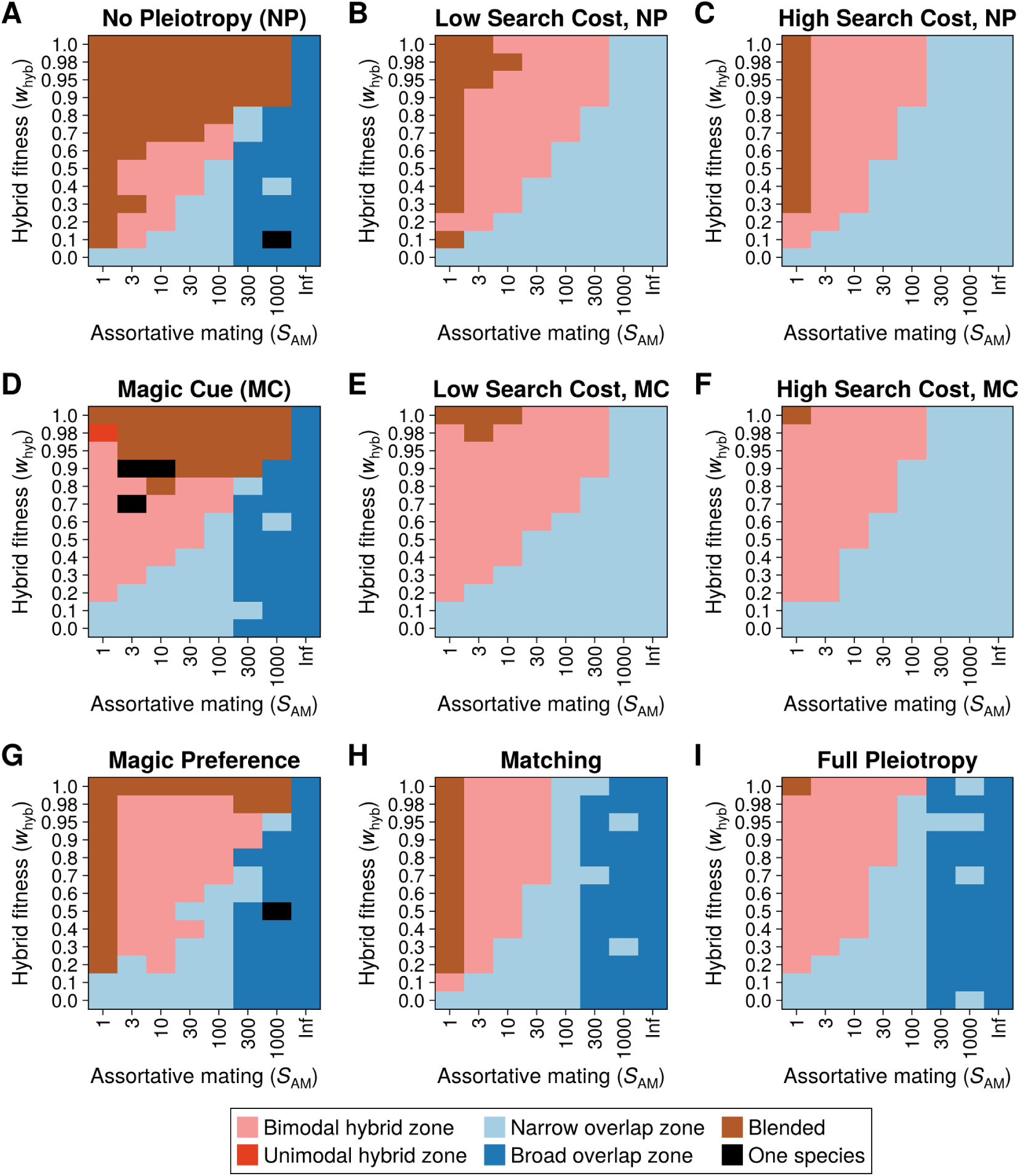
Most frequent outcomes for simulations conducted under identical conditions to Figure 3, but with one locus per trait instead of three. Here, search costs are very effective at maintaining species boundaries with or without a magic cue.

**FIGURE A-2.**
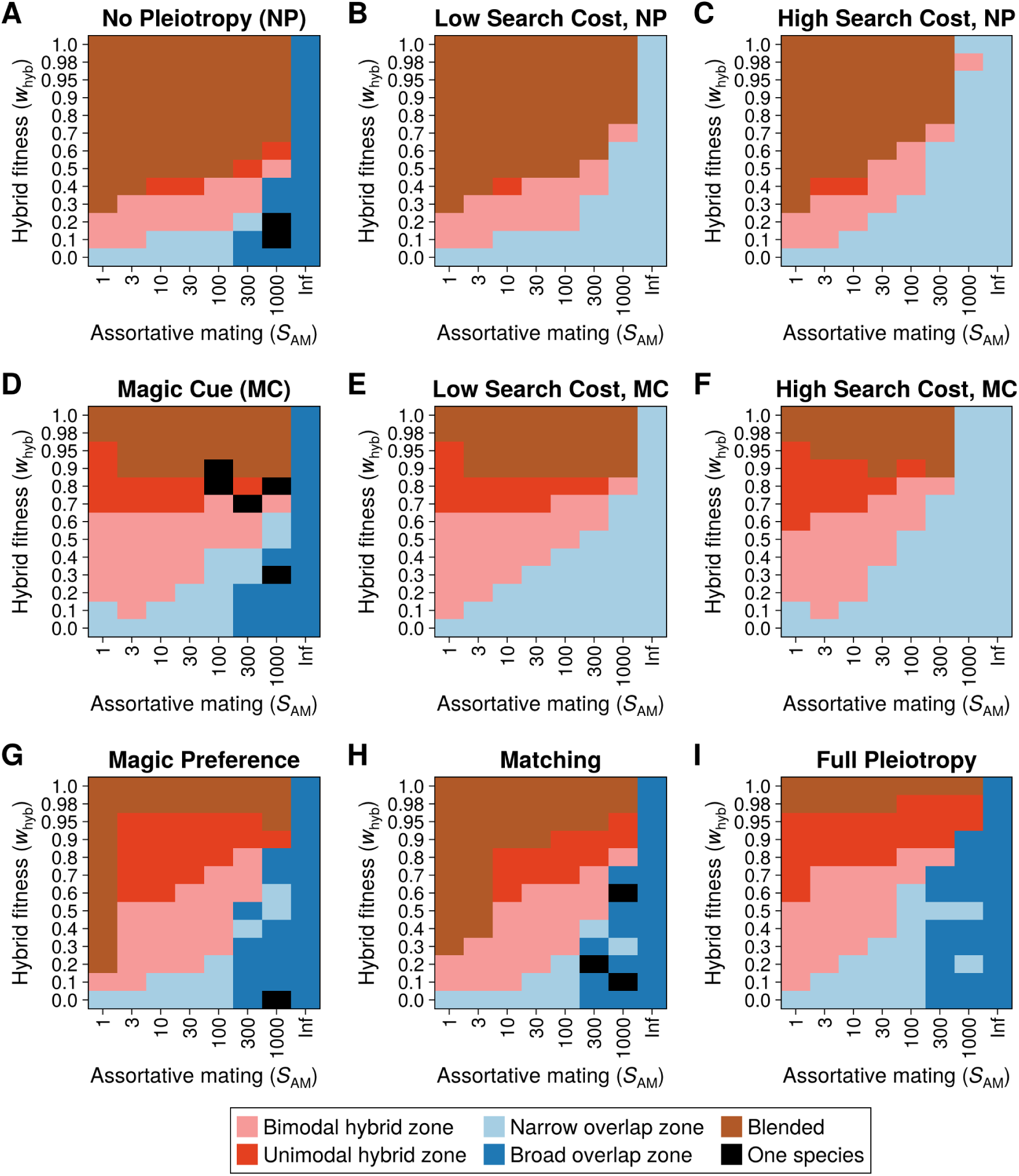
Most frequent outcomes for simulations conducted under identical conditions to Figure 3, but with nine loci per trait instead of three. Search costs are much less effective at maintaining stable clines than with one or three loci, but the magic cue results are similar.

**FIGURE A-3.**
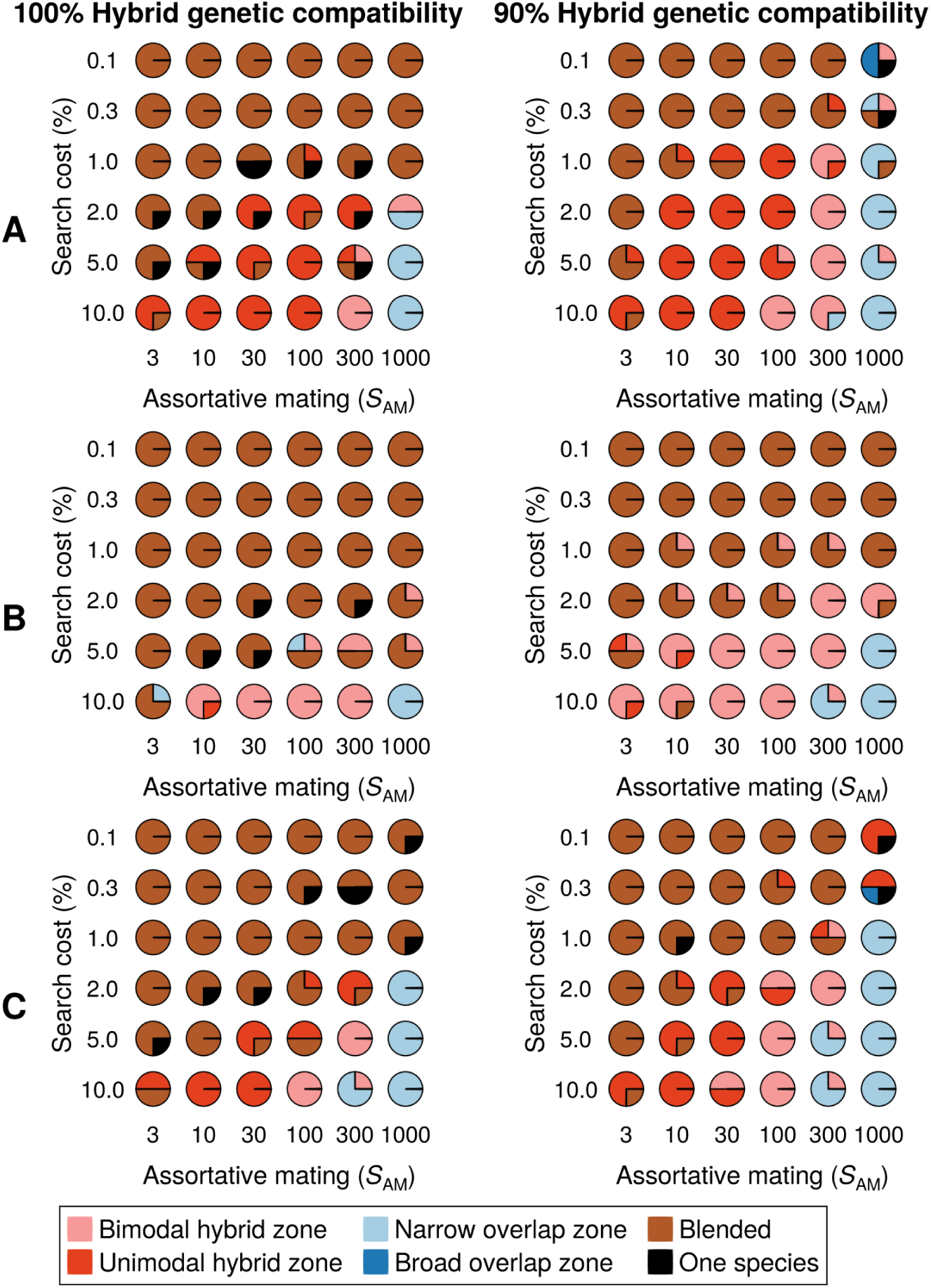
Outcomes of sets of four simulations conducted with varying search costs and strengths of conspecific mating for three different genetic architectures. A) One preference locus and three cue loci result in stable unimodal hybrid zones under a broad range of conditions. B) One cue locus and three preference loci are less stable, but can still maintain some bimodal hybrid zones. C) A multivariate preference trait produces stable hybrid zones similar to A). With the multivariate preference, three preference loci are paired with three cue loci so that the effects of each preference locus on match strength are independent.

## Notes

https://github.com/jrfarley/HZAM-2D/releases/tag/v2.0.1-review

### Competing Interest Statement

The authors have declared no competing interest.

